# Explaining Deep Learning-Based Representations of Resting State Functional Connectivity Data: Focusing on Interpreting Nonlinear Patterns in Autism Spectrum Disorder

**DOI:** 10.1101/2023.09.13.557591

**Authors:** Young-geun Kim, Orren Ravid, Xinyuan Zhang, Yoojean Kim, Yuval Neria, Seonjoo Lee, Xiaofu He, Xi Zhu

**Author notes:** Address correspondence to Xi Zhu, PhD, New York State Psychiatric Institute, Unit 69, 1051 Riverside Drive, New York, NY 10032. Equal contribution of the first two authors.

## Abstract

**Background:** Resting state Functional Magnetic Resonance Imaging fMRI (rs-fMRI) has been used to study brain function in psychiatric disorders, yielding insight into brain organization. However, the high dimensionality of the rs-fMRI data presents challenges, and requires dimensionality reduction before applying machine learning techniques. Neural networks, specifically variational autoencoders (VAEs), have been instrumental in extracting low-dimensional latent representations of resting state functional connectivity patterns, addressing the complex nonlinear structure of rs-fMRI. However, interpreting those latent representations remains a challenge. This paper aims to address this gap by creating explainable VAE models and testing their utility using rs-fMRI data in autism spectrum disorder (ASD).

**Methods:** One-thousand one hundred and fifty participants (601 HC and 549 patients with ASD) were included in the analysis. We extracted functional connectivity correlation matrices from the preprocessed rs-fMRI data using Power atlas with 264 ROIs. Then VAEs were trained in an unsupervised fashion. Lastly, we introduce our latent contribution scores to explain the relationship between estimated representations and the original rs-fMRI brain measures.

**Results:** We quantified the latent contribution scores for the ASD and control groups at the network level. We found that both ASD and control groups share the top network connectivity that contribute to all estimated latent components. For example, latent 0 was driven by resting state functional connectivity patterns (rsFC) within ventral attention network in both the ASD and control. However, significant differences in the latent contribution scores between the ASD and control groups were discovered within the ventral attention network in latent 0 and the sensory/somatomotor network in latent 2.

**Conclusion:** This study introduced latent contribution scores to interpret nonlinear patterns identified by VAEs. These scores effectively capture changes in each observed rsFC features as estimated latent representation changes, enabling an explainable deep learning model to better understand the underlying neural mechanism of ASD.

## Introduction

The use of functional magnetic resonance imaging (fMRI) data for the purpose of studying brain function, evaluating underlying neural mechanisms of treatment interventions, and aiding in the diagnosis of a variety of mental disorders has been around in psychiatry fields for the last few decades. In particular, resting state fMRI (rs-fMRI), measuring spontaneous neural activities in the absence of any specific task, has been used to reveal intrinsic patterns of brain functional connectivity and provide insights into brain functional organization. One of the most common ways to extract measures of brain networks of rs-fMRI data is to calculate the correlation coefficient between pairs of different regions of interest (ROIs) in the brain. The differences in connectivity between healthy subjects and those with psychiatric disorders have been examined to understand the underlying neural network deficits of those disorders (1). In recent years, machine learning approaches have gained prominence for the analysis of rs-fMRI for the diagnosis of psychiatric disorders and the prediction of treatment outcomes at the individual level (2).

Even so, many challenges still exist when trying to leverage rs-fMRI for these purposes. One of the main issues lies in the high dimensionality of the data. Very often, the rs-fMRI data can be partitioned into several hundred ROIs based on different brain atlases (3), e.g., 264 ROIs for the Power atlas (4) or 333 for the Gordon atlas (5). Resulting in connectivity matrices with more than ten thousand of image features. This creates challenges when using many standard machine learning techniques such as random forest, support vector machine, and regressions. Thus, dimensionality reduction methods are often used as a preprocessing step before applying machine learning algorithms to the rs-fMRI datasets. Traditional linear methods, including principal component analysis (PCA) and independent component analysis (ICA) have been used to transform the high-dimensional connectome features into a lower-dimensional space. However, these methods lack to address the complex nonlinear structure of rs-fMRI. In contrast, recently, many deep learning-based approaches have extracted low-dimensional latent factors (called *representation*) of resting state functional connectivity patterns (rsFC), showing remarkable performances with expressive nonlinear neural networks (6, 7).

As such, the advent of neural networks has provided a new avenue for dimensionality reduction. One model architecture which has gained prominence in the field of neuroimaging is the autoencoder (AE) framework (8). These models aim to learn a relatively low-dimensional latent representation of the original data, which can then be decoded to recover that data through the decoding phase. Additionally, to produce more effective and interpretable latent representations, variational autoencoder (VAE) approaches have been added and yielded promising results (9). Compared to AEs, VAE approaches possess prominent properties. 1) VAEs are probabilistic models that learn distributions of latent representations allowing for better modeling of complex data structures. They are an extension of nonlinear ICA (10), modeling the data generation mechanism with nonlinear neural networks and low-dimensional latent representations consisting of statistically independent components (11). In the implementation, representations follow user-specified distributions called *prior*, and one common choice is multivariate Gaussian distributions. 2) Compared to AEs, the training for VAEs is regularized to avoid overfitting and enforces the independency between components in estimated representations (12). Estimated representations are more interpretable when each component has a distinct role from all the others (13). For example, when analyzing hand-written digit images (e.g., MNIST http://yann.lecun.com/exdb/mnist/), VAEs may estimate two-dimensional representations where the first component explains the size of digits and the second one explains the slant. In the context of rs-fMRI data, certain latent components may explain within-network connectivity, while others explain between-network connectivity. 3) Moreover, VAEs provide personalized inference on the latent space. They approximate the distribution of latent representations given observations, allowing for subject-level information such as uncertainty or variance of estimated representations.

Despite the improvement in the performance of machine learning models when using these latent features, interpreting the significance of the latent features is a challenge. Many neural network models can, at first glance, appear to be black boxes with complex and difficult-to-interpret representations. However, across the deep learning community, strides have been made to create tools for interpreting these models. Initially, much of the work in visualizing the latent features of autoencoder models was done in the field of computer vision. The tools developed there benefited from the fact that images are relatively easy to interpret and understand naturally by humans. Thus, it was easier to see the qualitative contribution of each latent feature as you could directly examine the images produced by these tools. However, brain imaging modalities such as rs-fMRI do not share the same natural interpretability.

Some initial efforts have been made to generate interpretable latent representations of rs-fMRI data using VAEs. Kim et al., trained VAEs with large rs-fMRI data from Human Connectome Project. 2D grids of rs-fMRI patterns at every time point were extracted as an image and input to the VAE models. The results demonstrated that estimated representations from VAEs effectively characterized individual identification (14). Another study applied VAEs to rs-fMRI data from Autism Brain Imaging Data Exchange (ABIDE) and found an autism spectrum disorder (ASD)-related latent factor (15). However, the dimension of representations was two, and this study lacks explanation of the complicated information in rs-fMRI related to ASD. Moreover, latent representations were formed by individual brain regions (e.g., frontal cortices and frontoparietal) rather than brain networks (e.g., ECN, SN, and DMN).

The goal of this paper is to extract the latent representation of VAE models trained on rs-fMRI data, and create explainable VAE models by visualizing and quantifying the latent representation based on the input rs-fMRI brain features. We will test the utility of this tool using the rs-fMRI dataset.

## 1. Methods

### Dataset

We used publicly available data from Paris-Saclay Center for Data Science that was initially published for competition in the Imaging-Psychiatry Challenge (IMPAC; https://paris-saclay-cds.github.io/autism_challenge/). One-thousand one hundred and fifty participants (601 HC, 549 patients with ASD) from 35 sites were included in the analysis (see Table 1 for demographics characteristics). We further removed 121 participants that failed to pass quality control procedures. The IMPAC dataset includes T1 structural, rs-fMRI images, data acquisition site, age, and sex.

**Table 1:** The descriptive information for each site from the public dataset.

| Site | Total N | Female | Male | Mean Age | StdAge | MinAge | Max Age | Ctrl | ASD | Reject | Accept |
| --- | --- | --- | --- | --- | --- | --- | --- | --- | --- | --- | --- |
| 0 | 45 | 0 | 45 | 39.96 | 14.16 | 18 | 62 | 23 | 22 | 6 | 39 |
| 1 | 32 | 7 | 25 | 7.93 | 0.97 | 6 | 11 | 15 | 17 | 13 | 19 |
| 2 | 35 | 11 | 24 | 10.43 | 1.73 | 8 | 14 | 21 | 14 | 1 | 34 |
| 3 | 32 | 18 | 14 | 23.13 | 11.83 | 8 | 47 | 23 | 9 | 5 | 27 |
| 4 | 17 | 6 | 11 | 26.76 | 10.01 | 17 | 54 | 11 | 6 | 1 | 16 |
| 5 | 102 | 38 | 64 | 10.43 | 1.25 | 8 | 13 | 76 | 26 | 5 | 97 |
| 6 | 21 | 0 | 21 | 23.38 | 3.81 | 18 | 33 | 0 | 21 | 0 | 21 |
| 7 | 48 | 5 | 43 | 10.11 | 5.98 | 5 | 35 | 16 | 32 | 0 | 48 |
| 8 | 18 | 3 | 15 | 6.69 | 0.98 | 5 | 9 | 0 | 18 | 6 | 12 |
| 9 | 55 | 22 | 33 | 11.47 | 2.03 | 8 | 15 | 29 | 26 | 1 | 54 |
| 10 | 32 | 6 | 26 | 13.30 | 2.97 | 7 | 18 | 13 | 19 | 1 | 31 |
| 11 | 23 | 0 | 23 | 15.43 | 2.79 | 12 | 20 | 12 | 11 | 3 | 20 |
| 12 | 20 | 5 | 15 | 15.13 | 1.70 | 12 | 18 | 8 | 12 | 2 | 18 |
| 13 | 20 | 4 | 16 | 22.12 | 7.67 | 11 | 39 | 10 | 10 | 2 | 18 |
| 14 | 23 | 6 | 17 | 30.86 | 12.07 | 18 | 56 | 11 | 12 | 4 | 19 |
| 15 | 16 | 5 | 11 | 27.31 | 6.42 | 19 | 40 | 6 | 10 | 3 | 13 |
| 16 | 32 | 8 | 24 | 10.38 | 1.31 | 8 | 13 | 18 | 14 | 2 | 30 |
| 17 | 18 | 0 | 18 | 21.89 | 2.66 | 18 | 29 | 10 | 8 | 0 | 18 |
| 18 | 19 | 5 | 14 | 14.31 | 1.30 | 12 | 17 | 12 | 7 | 3 | 16 |
| 19 | 40 | 6 | 34 | 28.60 | 11.76 | 7 | 58 | 26 | 14 | 7 | 33 |
| 20 | 106 | 25 | 81 | 15.52 | 7.01 | 7 | 39 | 61 | 45 | 5 | 101 |
| 21 | 11 | 0 | 11 | 10.59 | 1.38 | 8 | 13 | 8 | 3 | 0 | 11 |
| 22 | 24 | 3 | 21 | 17.00 | 3.57 | 10 | 24 | 12 | 12 | 3 | 21 |
| 23 | 38 | 6 | 32 | 20.03 | 7.13 | 9 | 35 | 17 | 21 | 6 | 32 |
| 25 | 19 | 0 | 19 | 37.37 | 8.37 | 27 | 64 | 9 | 10 | 7 | 12 |
| 26 | 23 | 6 | 17 | 14.54 | 2.01 | 9 | 17 | 15 | 8 | 1 | 22 |
| 27 | 19 | 3 | 16 | 10.13 | 1.55 | 8 | 12 | 9 | 10 | 3 | 16 |
| 28 | 32 | 0 | 32 | 17.32 | 3.83 | 12 | 26 | 14 | 18 | 2 | 30 |
| 29 | 38 | 6 | 32 | 12.84 | 2.23 | 8 | 18 | 18 | 20 | 4 | 34 |
| 30 | 15 | 1 | 14 | 12.42 | 1.21 | 10 | 15 | 6 | 9 | 4 | 11 |
| 31 | 59 | 17 | 42 | 13.38 | 2.85 | 8 | 19 | 37 | 22 | 9 | 50 |
| 32 | 28 | 2 | 26 | 15.84 | 3.60 | 13 | 29 | 16 | 12 | 2 | 26 |
| 33 | 62 | 0 | 62 | 22.10 | 8.01 | 9 | 50 | 27 | 35 | 6 | 56 |
| 34 | 28 | 6 | 22 | 12.90 | 2.98 | 7 | 18 | 12 | 16 | 4 | 24 |
| sum | 1150 | 230 | 920 |  |  |  |  | 601 | 549 | 121 | 1029 |

### Image acquisition and processing

All time-series imaging data were acquired using specific atlases and a fetcher provided by IMPAC. We extracted functional connectivity matrices from the preprocessed rs-fMRI data using Power atlas with 264 ROIs (4). The ROI names and the networks to which they belong are defined at https://www.jonathanpower.net/2011-neuron-bigbrain.html. Pairwise correlations were calculated between each ROI for each participant and transformed into a correlation matrix whose elements are Pearson correlations ranging from –1 to 1. Finally, we vectorized correlation matrices by flattening the lower triangle part. For a connectivity matrix with N ROIs, the length of the 1D vectorized correlation vector was calculated by (N – 1) x N/2. The 1D vector was used as an input signal in VAE models. To correct site effects and adjust for age and gender covariates, we performed a Combat algorithm on the correlation vectors using neuroHarmonize (16).

### Variational autoencoders

#### Model Architectures

We denote high-dimensional observations and low-dimensional representations by *X* ∈ ℝ^*m*^ and *Z* ∈ ℝ^*n*^, respectively. Realizations of random variables are denoted by small characters. VAEs consist of two parts: (i) encoders modeling the distribution of representations given observations, *q*_*ϕ*_(*z*|*x*), and (ii) decoders modeling the distribution of observations given representations, *p*_*θ*_(*x*|*z*) (Figure 1), where *ϕ* and *θ* are neural network parameters. Both *q*_*ϕ*_(*z*|*x*) and *p*_*θ*_(*x*|*z*) are usually modeled as multivariate Gaussian distributions with diagonal variances, *N*(*μ*_*ϕ*_(*z*|*x*), Σ_*ϕ*_(*z*|*x*) and *N*(*μ*_*θ*_(*x*|*z*),*I*_*m*_), respectively, to apply reparameterization trick (9) where *I*_*m*_ is the identity matrix of size *m*. Encoders extract representations from observations, decoders reconstruct the original data with them, and they are trained by maximizing evidence lower bounds (ELBOs) (17). For example, for a given *x*, we can first sample *z* following *N*(*μ* _*ϕ*_(*z*|*x*), Σ_*ϕ*_(*z*|*x*)), and then use *μ*_*θ*_ (*x*|*z*) as reconstruction results.

**Figure 1:**
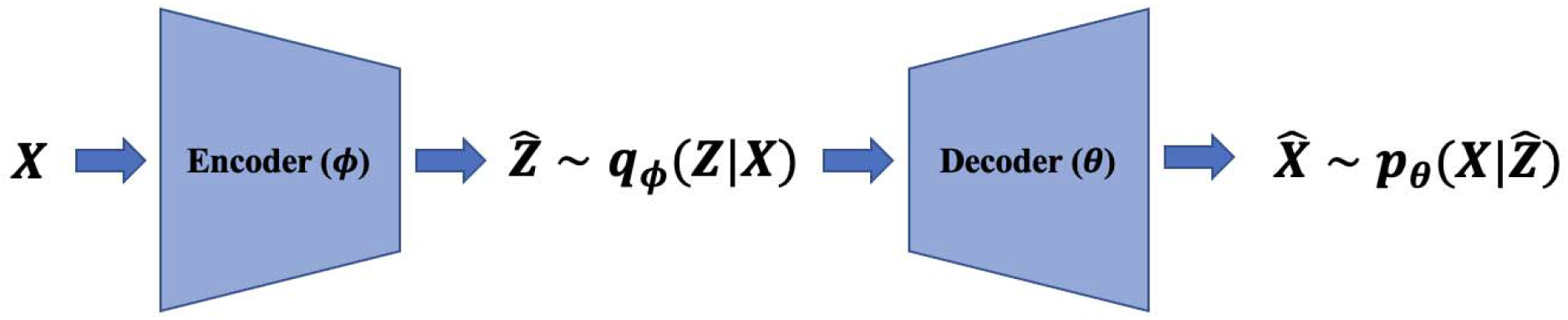
VAEs model the data generation mechanism with low-dimensional representations and neural networks called decoders. The encoders estimate representations with observations and decoders reconstruct the original data with representations.

Compared to AEs, VAEs are distinct in that they are generative models, i.e., they model distributions of observations. Latent representations *Z* consist of statistically independent components and nonlinear decoders input *Z* to model, *p*_*θ*_(*x*|*z*), *I*_*m*_) e.g., *N*(*μ*_*θ*_ (*x*|*z*),*I*_*m*_) When we assume that the *p*_*θ*_(*x*|*z*) has zero-variance, the data generation structure of VAE reduces to nonlinear ICA (18). By estimating the nonlinear decoders with likelihood maximization and approximating their inverse mapping with encoders, VAEs can separate blind sources from high-dimensional complicated observations, e.g., functional connectivity.

#### Loss Function

The loss function of VAEs is negative ELBO, 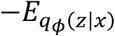 log *p*_*θ*_(*x*|*z*)+*D*_*KL*_(*q*_*ϕ*_(*z*|*x*) ‖*p*(*z*)), and is an upper bound of the negative data log-likelihood, - log *p*_*θ*_ (*x*), where *D*_*KL*_ denotes the Kullback-Leibler (KL) Divergence and *p*(*z*)denotes a user-specified prior distribution of representations, e.g., multivariate standard Gaussian distributions. Minimizing the loss function of VAEs is equivalent to maximizing a lower bound of likelihoods. In the loss function, the first term is called reconstruction error which measures how reconstruction results differ from the original observations, and it is the mean squared error when we use Gaussian distributions for decoder distributions. The second term measures the discrepancy between *q*_*ϕ*_(*z*|*x*) and *p*(*z*). Considering that the loss function of AEs is the reconstruction error, training VAEs can be viewed as training AEs with the KL-regularization term for the learning data generation mechanism.

### Latent contribution scores to explain nonlinear representations

#### Latent Contribution Scores

We introduce our latent contribution scores to explain the relationship between estimated representations and the original rs-fMRI brain measures.

For any observation *x* ∈ ℝ^*m*^, encoder parameter *ϕ*, and decoder parameter *θ*, to measure the contribution of each component in *z* ∈ ℝ^*n*^, we propose a matrix *D*(*x*) whose elements are latent contribution scores,

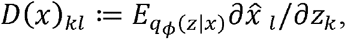

where 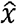 denotes the reconstruction result, *k* =1, …, *n*, and *l* =1, …, *m*.

The proposed latent contribution scores are interpretable in two respects: (i) they are extensions of mixing weights in ICA to nonlinear data generation mechanism, and (ii) they are input perturbation-based scores (19, 20). For (i), the 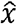 is the reconstruction result with estimated sources whose gradients are mixing weights under a linear generation mechanism. Similar to the explanation on mixing weights in ICA, we can explain the contribution of estimated representations on reconstructions as follows: The increment of *k* -th element of estimated representations by one unit yields the increment of the *l*-th element of reconstructed observations, e.g., *l*-th element of reconstructed rs-fMRI functional connectivity, by *D*(*x*)_*kl*_ unit on average over *q*_*ϕ*_(*z*|*x*). With these scores, we can explain how estimated representations change reconstructions in each group, e.g., ASD and healthy control groups. For (ii), in interpretable machine learning literature, input perturbation-based scores are a type of feature importance measures used to explain how outputs of complicated and nonlinear networks react to the perturbation on latent components. This provides an estimate of how important each feature is for the model’s decision-making process. Our scores are input perturbation-based scores in which they average gradients, the marginal changes of outputs by decoders with respect to input (estimated) representations.

#### Numerical Approximation for Latent Contribution Scores

Numerical approximation for the proposed latent contribution scores consists of two parts: (i) approximating gradients for a given representation and (ii) averaging gradients computed in (i) over encoder distributions. For (i), we computed average slopes with small perturbations. Let *x* be an observation and 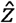 be an estimated representation sampled from *q*_*ϕ*_(*z*|*x*). We first compute reconstructions using 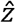 and 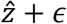, denoted by 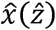 and 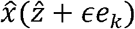, respectively, where *e*_*k*_ denotes the *k*-th component of standard basis of ℝ^*n*^, and then compute 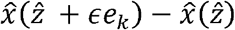 which is a numerical approximation of partial gradients up to constant multiplication. For (ii), in computing scores for the *k*-th latent component, we used fixed points rather than sampling to provide deterministic scores. For all axes except the *k*-th axis, we used mean of encoder distributions, and for the *k* -th axis, we used pre-specified grid points ranging from mean minus three standard deviations to plus three standard deviations. We first approximate gradients at each grid point, and then average them to compute latent contribution scores.

### Explaining rs-fMRI and Brain Networks with Latent Contribution Scores

#### Model architecture

In our experiments on the rs-fMRI dataset, the observations *X* are the lower triangle part of functional connectivity matrices from the Power atlas with 264 ROIs. Each element is the correlation of the resting state activity between one of 264 brain regions and another (Figure 2). Both the encoder and decoder have one hidden layer. The sizes of the respective layers were chosen by performing a sparse grid search for each of the layers’ sizes independently and evaluating the performance of the model both with respect to the loss function (21). For hyperparameter tuning, we considered the following choices: {tanh, SELU} for activation functions (22); {20, 40, 50, 80, 100, 150, 200, 250} for the number of the hidden node; {2, 5, 10, 15, 20} for the latent dimension. We used the loss of VAEs for the model selection criterion. Both the encoder and decoder have one hidden layer. The sizes of the respective layers were chosen by performing a sparse grid search for each of the layers’ sizes independently and evaluating the performance of the model, both with respect to the loss function (21). The chosen number of hidden nodes and latent dimension are 80 and 5, respectively.

**Figure 2:**
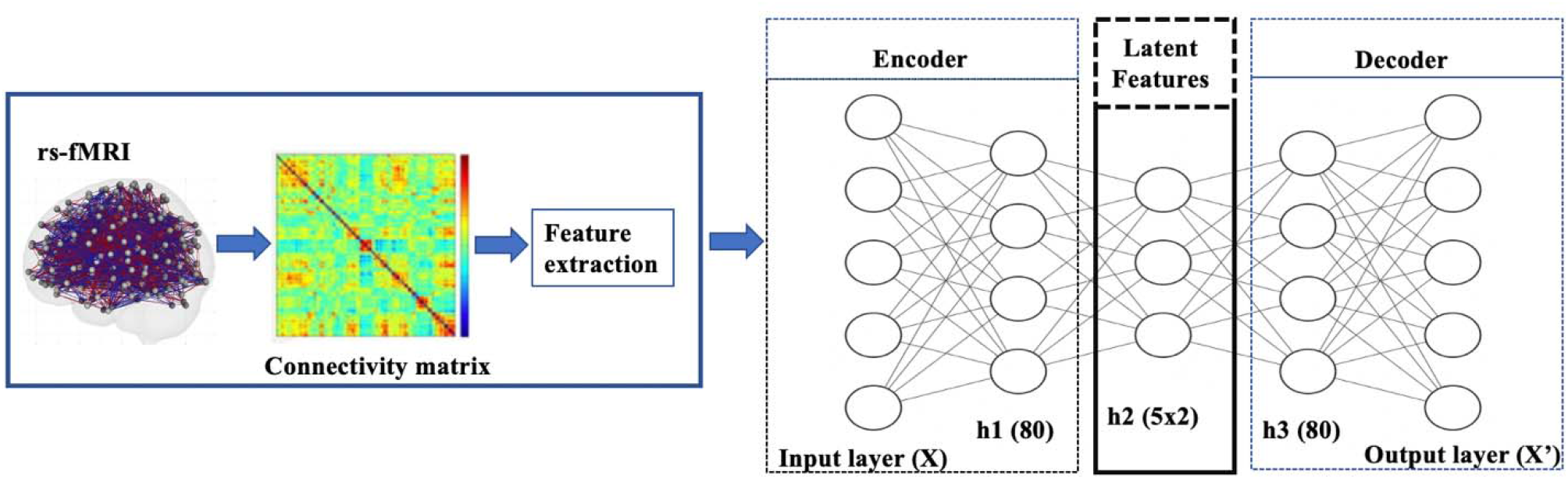
Diagram of VAE pipeline: The model was trained using rs-fMRI data. The samples were then split into a training+validation (70%) and independent-test (30%) data. Then 20% of the training data was set aside for validation and hyperparameter tuning. Once the training+validation was completed, the model’s performance was evaluated on the independent-test data, which provides an unbiased estimate of how the model generalizes to unseen data. The resulting VAE model learned to encode patterns from the input brain features into its latent representation.

#### Training of VAEs

We trained VAEs in an unsupervised fashion without using labels about ASD and control groups. We standardized data with median and interquartile ranges and added Gaussian noise with a zero-mean and standard deviation of 0.1 to input data for denoising purposes to learn robustness representations (23). The whole dataset was split into training+validation (70%), and test (30%) sets. The batch size, the number of epochs, and the weight decay was 128, 1000, and 0.1, respectively. We applied *L*_2_ regularization. For stopping criterion to evaluate convergence, we used the validation loss, negative ELBO.

## Results

We compared latent contribution scores of each latent variable based on the rsFC. Figure 3 provides a visualization of the top 0.05% resting-state functional connectivity with the highest latent contribution scores. The depicted brain network features change the most as the estimated representation (latent) changes.

**Figure 3:**
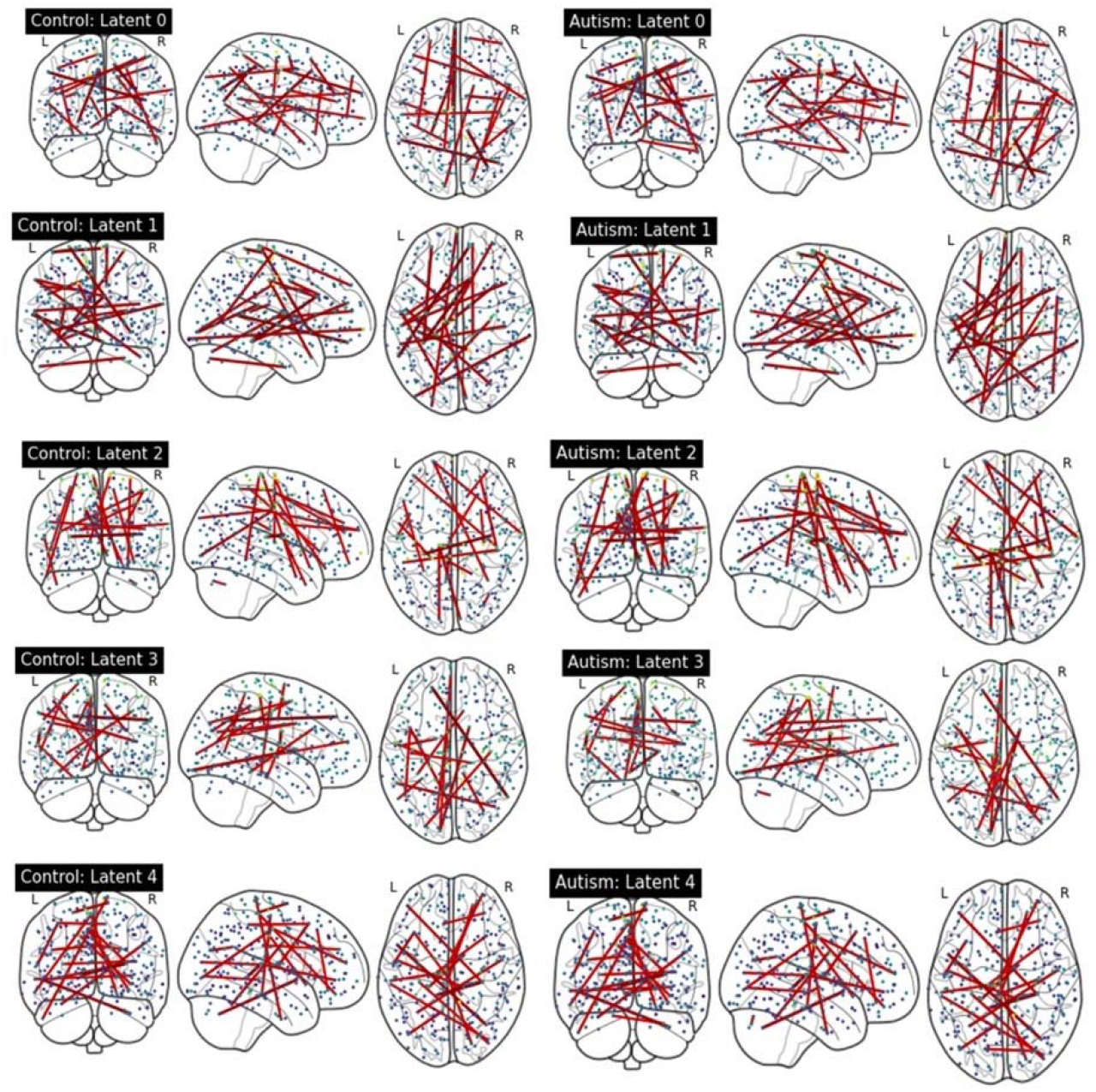
Visualization of top 0.05% functional connectivity based on latent contribution scores. Each row shows results on each component of estimated representations. The left and right columns display results on the control and autism groups, respectively.

We quantified the latent contribution scores for the ASD and control groups at the network level, as detailed in Table 2 and Figure 4. The ASD and control groups share the top network connectivity for all estimated latent components. For example, within ventral attention network (VAN) contributes the most to latent 0 in both the ASD and control. Similarly, latent 1 is primarily influenced by the rsFC between somatomotor (SMN)-memory retrieval networks, latent 2 is driven by the rsFC within SMN; latent 3 is driven by rsFC between memory retrieval-cerebellar networks, and latent 4 is driven by rsFC between cerebellar-dorsal attention networks in both groups.

**Table 2:** Summary of top 3 network connectivity having the highest latent contribution scores. We averaged scores from rsFC across ROIs over the network level.

| **Latent Representation** | **Latent contribution score rank** | **Control Group** | **Autism Group** |
| --- | --- | --- | --- |
| Latent 0 | **Top 1** | **Within ventral attention** | **Within ventral attention** |
|  | **Top 2** | **Between memory retrieval and subcortical** | **Between memory retrieval and ventral attention** |
|  | **Top 3** | **Within subcortical** | **Within uncertain** |
| Latent 1 | **Top 1** | **Between sensory/somatomotor mouth and memory retrieval** | **Between sensory/somatomotor mouth and memory retrieval** |
|  | **Top 2** | **Within sensory/somatomotor mouth** | **Within sensory/somatomotor mouth** |
|  | **Top 3** | **Within uncertain** | **Within uncertain** |
| Latent 2 | **Top 1** | **Between memory retrieval and ventral attention** | **Within sensory/somatomotor hand** |
|  | **Top 2** | **Within sensory/somatomotor hand** | **Between sensory/somatomotor hand and cerebellar** |
|  | **Top 3** | **Between sensory/somatomotor hand and memory retrieval** | **Between sensory/somatomotor hand and memory retrieval** |
| Latent 3 | **Top 1** | **Between memory retrieval and cerebellar** | **Between memory retrieval and cerebellar** |
|  | **Top 2** | **Within cerebellar** | **Within sensory/somatomotor hand** |
|  | **Top 3** | **Within sensory/somatomotor hand** | **Between sensory/somatomotor hand and memory retrieval** |
| Latent 4 | **Top 1** | **Between cerebellar and dorsal attention** | **Between cerebellar and dorsal attention** |
|  | **Top 2** | **Within dorsal attention** | **Within dorsal attention** |
|  | **Top 3** | **Within ventral attention** | **Between uncertain and memory retrieval** |

**Figure 4:**
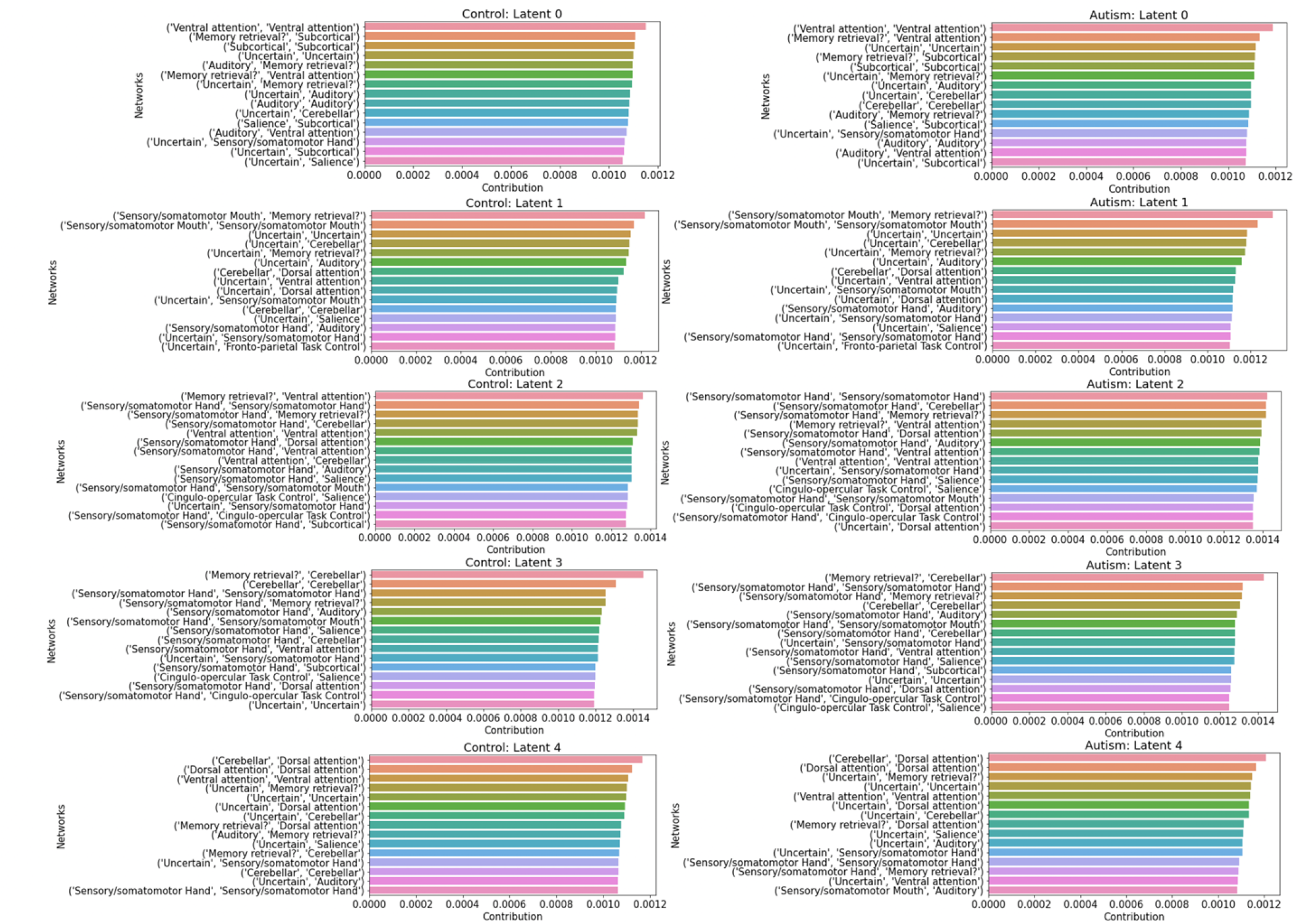
Summary of top 15 network connectivity having the highest latent contribution scores. We averaged scores from rsFC across ROIs over the network level.

Among the rsFC network ROIs that contribute the most to each latent component (as shown in Table 2), we further compared them between the ASD and control groups. We first conducted t-test to filter ROIs with significantly different latent contribution scores between ASD and control groups, using a significance level of 1%. Subsequently, we averaged the scores at the network level. We found significant difference of the latent contribution scores in the VAN network (top network that driven the latent 0), and the SMN (top 2 network that driven the latent 1).

## Discussion

In this study, we proposed latent contribution scores to explain nonlinear patterns identified by VAEs. These proposed scores effectively capture marginal changes of each component of observations as estimates representations change. With this toolkit, we were able to examine both quantitative and qualitative differences in a VAE-based model’s representation of psychiatric disorders.

Specifically, we were able to quantify which brain networks were most significantly contribute to each latent component that differentiated between ASD and controls. Five latent components were identified including within VAN driven latent 0; SMN-memory retrieval networks driven latent 1, memory retrieval-cerebellar networks driven latent 3, and cerebellar-dorsal attention networks driven latent 4. Among these 5 latent components, the latent contribution scores in the ventral attention network driven latent 0, and the SMN - driven latent 1 are significantly different between the ASD and control groups. The VAN and SMN are two important networks implicated in ASD. VAN plays a crucial role in processing sensory information and direct attention. Studies suggest that individual with ASD have altered rsFC in the VAN, which could contribute to difficulties with focusing, maintaining and shifting attention and social communication (24). The SMN is involved in the processing of sensory information and controlling motor functions. Individuals with ASD often show different sensory processing compared with controls. Altered rsFC in the SMN could potentially contribute to these sensory processing differences. Moreover, altered rsFC in SMN may contribute to the motor coordination difficulties in individuals with ASD (25).

Our approach is generally applicable to a broader class of dimension reduction methods, including autoencoder and its derivatives, bidirectional generative adversarial networks (26, 27), and deep belief networks (28) that use probabilistic encoders and any desired imaging modality. In fact, our technique does not require the model to even be a neural network as long as it has mapping from high dimentional observations to estimated representations and vice versa, and the gradient can be numerically approximated. Some examples of models that fall in this category are VAE-based GANs (29) and Hyperspherical VAE (30). Another advantage of our approach lies in its visualization capability. When observations are visually perceived as in natural images, we can display our latent contribution scores. For example, when the data modality is 4D fMRI voxel-time space data, we can visualize the proposed contribution scores for each latent component on the 4D space and interpret their spatial-temporal patterns.

It is important to note that our analysis was only done on a VAE applied to ROI-to-ROI measures extracted from RS timeseries data. Other RS measurements such as ALFF and ReHo have not used for this analysis and we may try in the future work. Additionally, we used the Power atlas (Power et al. 2011) with 264 ROIs in this study. Future studies could try to leverage different functional connectivity matrices using different atlases as well. Moreover, since different MRI modalities contain complementary information for ASD, such as task-based fMRI, T1 structural MRI or diffusion weighted imaging, fusing multiple modalities may provide additional information, and contribute to each latent component. The proposed deep learning model can potentially be used to combine different imaging modalities via stacked autoencoder, and explain the contribution of each modality to the latent components, which can help to understand the mechanism of psychiatric disorders such as ASD.

In conclusion, our proposed latent contribution scores provide a degree of interpretability of deep learning models. The models that applied to the rs-fMRI data can be understood and interpreted by humans. Moreover, explainable VAEs can provide insights into what features from single modality or a combination of multiple modalities are most important for a particular prediction task (such as classification of ASD from controls), this can be valuable for feature engineering and understanding the underlying neural mechanisms.

## Acknowledgements

Dr. Zhu is supported by NIH K01MH122774 and by a NARSAD Young Investigator Grant from the Brain & Behavior Research Foundation 27040; Dr. Kim is supported by NIH R01MH124106.

## Code availability

https://github.com/kyg0910/Deep-Learning-Based-Representations-of-Resting-State-Functional-Connectivity-Data

